# Cancer evolution simulation identifies possible principles underlying intratumor heterogeneity

**DOI:** 10.1101/022806

**Authors:** Atsushi Niida, Satoshi Ito, Georg Tremmel, Seiya Imoto, Ryutaro Uchi, Yusuke Takahashi, Koshi Mimori, Satoru Miyano

## Abstract

Cancer arises from accumulation of somatic mutations and accompanying evolutionary selection for growth advantage. During the evolutionary process, an ancestor clone branches into multiple clones, yielding intratumor heterogeneity. However, principles underlying intratumor heterogeneity have been poorly understood. Here, to explore the principles, we built a cellular automaton model, termed the BEP model, which can reproduce the branching cancer evolution in silico. We then extensively searched for conditions leading to high intratumor heterogeneity by performing simulations with various parameter settings on a supercomputer. Our result suggests that multiple driver genes of moderate strength can shape subclonal structures by positive natural selection. Moreover, we found that high mutation rate and a stem cell hierarchy can contribute to extremely high intratumor heterogeneity, which is characterized by fractal patterns, through neutral evolution. Collectively, This study identified the possible principles underlying intratumor heterogeneity, which provide novel insights into the origin of cancer robustness and evolvability.

## Introduction

Cancer is a heterogeneous disease. Its molecular characteristics differ among patients, who therefore require personalized therapies. In addition to this intertumor heterogeneity, another level of heterogeneity, called intratumor heterogeneity, exists; within even in a single tumor, genetically distinct clones co-exist. Understanding intratumor heterogeneity is clinically important, since high intratumor heterogeneity presumably leads to therapy resistance, increasing a probability that a tumor harbors resistant clones. Intratumor heterogeneity can be shaped by branched evolution of the cancer genome [29, 51].

During evolution of the cancer genome. multiple driver mutations are accumulated, which give a selective advantage to clones through either increasing its survival or reproduction. We also observe many passenger mutations, which occur in the same genome with driver mutations, but have no effect on the fitness of clones. While driver mutations tend to cause clonal expansions, passenger mutations may be associated with a clonal expansion just by ‘hitchhiking’ driver mutations [43]. Classically, it has been assumed that multiple driver mutations are acquired in some specified order and cancer evolution has been viewed as a linear process where successive clonal expansions are caused by acquisition of each driver mutation.

On the other hand, many studies have also observed extensive intratumor heterogeneity, which cannot be explained by such a linear model. Recently, multiregional sequencing, which profiles cancer genomes obtained from geographically separated multiple regions in a single tumor, has revealed extensive inratumor heterogeneity for several types of solid tumors, and established that cancer evolution is a highly branched process [50, 16, 15, 44, 52, 8]. From multiregional sequencing data, a new categorization of mutations, which is different from the above-described drive/passenger categorization, has emerged: fonder and progressor mutations. Founder mutations are observed in all regions, while progressor mutations are not shared by all regions and shapes intratumor heterogeneity. For an evolutionary view, the two types of mutations are seen as follow. early in the evolution, founder mutations are accumulated to establish an single ancestor clone. the ancestor clone is then branched into multiple subclones while accumulating progressor mutations. Depending on studies, founder and progressor mutations are referred as to trunk and branch mutations, by positioning them on evolutionary trees. Also, multiregional sequencing studies suggest that the degree of intratumor heterogeneity is associated with the malignancy of cancer [52]; however, further studies are necessary to clarify this point.

Mathematical modeling have a long history in cancer research and is a powerful tool for studying cancer evolution [4]. For example, cancer evolution has been analytically studied as Moran process, Wright-Fisher process and branching process[11, 20, 36, 7, 10, 18, 3, 34]. However, those analytical approaches have limitations for build a realistic biological model: e.g., the limited numbers of states in branched evolution and assumption of constant population size. On the other hand, although analytical results are generally unavailable, a cellular automaton model (or referred as to a agent-based model) has the advantage that evolutionary rules are explicitly and flexibly modeled [12]. To date, various simulation studies employing cellular automaton models succeeded in reproducing intratumor heterogeneity [17, 40, 13, 28, 41, 42]. However, to our best knowledge, there exists no model that reproduces the recently emerging view of highly branched cancer evolution, and evolutionary principles underlying intratumor heterogeneity have not been fully explored.

In this study, we propose a new mathematical model of highly branched cancer evolution. Our branching evolutionary process (BEP) model is a cellular automaton model where an automaton represents a biological cell having dozens of mutable genes. A state in branched evolution is defined by the combination of mutated genes, which can produce the virtually infinite number of branched states. Also, some of the genes are assumed as driver genes whose mutation confer growth advantage to the cell, and simulation results are compatible with the driver/passenger mutation paradigm. Moreover, running thousands of simulations with different parameter settings on our supercomputer [32], we extensively searched for conditions producing high intratumor heterogeneity. Our BEP simulation successfully identified several possible evolutionary principles underlying intratumor heterogeneity.

## Result

### Summary of our methodology

To reproduce genomic intratumor heterogeneity *in silico,* we modeled branched evolution as the BEP model using a cellular automaton model (Figure 1). We assumed that each cell in a tumor acts as a cellular automaton. Each cell contains *n* genes, of which *d* genes are driver genes. At each time step, each cell dies with a probability *q* and divides into two daughter cells with a probability *p*. At each cell division, each gene is mutated with a probability *r*. A mutation in a driver gene is referred to as a diver mutation and, if a cell obtains one driver mutation, the division probability is increased by 10*^f^*-fold from the next time step, where the parameter *f* represents the strength of the driver genes. We also extended the model and analyzed effects of a stem cell hierarchy as described later.

**Figure 1:**
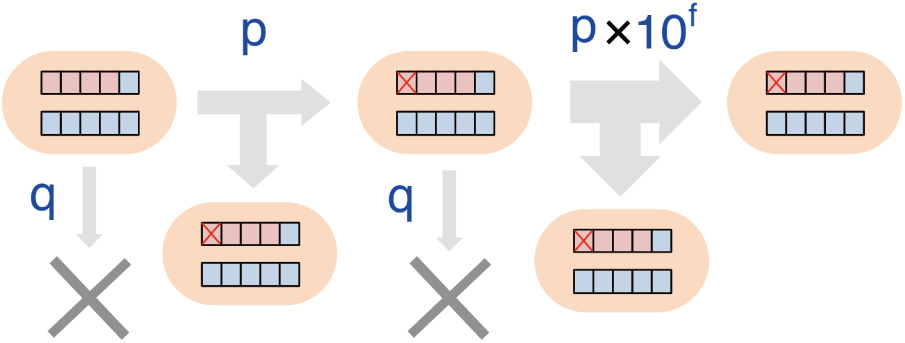
A schema of the BEP model. A cell has *n* genes, *d* out of which are driver genes. In this schema, *n* = 10 and *d* = 4, and red and blue boxes denote driver and non-drive genes, respectively. In a unit time step, a cell divides or dies with probabilities *p* and *q*, respectively. During each cell division, each gene is randomly mutated with a probability *r*, and one driver mutation, which is denoted by a red cross, increases *p* by 10*^f^*-fold.

Starting from *c*_0_ normal cells, having no mutation, we grew the tumor until the number of cells in the tumor (hereafter referred as to population size) reached *c_max_.* In our simulation, a set of mutated genes in a cell is referred to as a genotype, while a set of cells that have an identical genotype is referred to as a clone. To visualize tumor growth, we spread the dividing cells over a two-dimensional space and colored each clone in the tumor so that color similarity represents similarity between the genotypes of clones.

To assess intratumor heterogeneity, we randomly sampled *m* cells from the simulated tumor and visualized their genome as a mutation profile matrix. As a quantitative measure of intratumor heterogeneity, we calculated entropy statistic from a distance matrix among genotypes of the *m* cells, utilizing singular value decomposition [33]. We termed the statistic population entropy, which is represented by *ϵ*. Several existing measures are available for the same purpose [17, 40, 42]; however, they treat different clones equally. Since each pair of clones can be differently similar in our model, we employed a new measure that can consider genotype similarity among clones. In addition to *ϵ*, we calculated the number of founder mutations *ρ* to characterize mutation profiles. We also obtained a number of other statistics, including the average count of mutation per cell *μ*, population fitness *λ*, and the time step *τ* at which the tumor size reaches *C_max_*.

To search for principles underlying intratumor heterogeneity, we performed simulations with various parameter settings and examined parameter dependence of the statistics. In this study, we examined 2610 parameter settings in total. For each parameter settings, we repeated the simulation 20 times and calculated the average of each statistic. The simulations were performed in parallel on our supercomputer system [32], which enabled an extensive parameter search in a practical time. On average, each simulation took about 2.75 CPU core hours. In total, our study needed 2.75 × 2610 × 20 =143550 CPU core hours ≈ 16.4 CPU core years. However, by parallelizing thousands of simulations, they finished within a couple of days. Results from the simulations can be interactively explored at a supplementary website: http://bep.hgc.jp/.

### Effects of driver gene size and strength

From the results of the extensive parameter search, we found several conditions that yield high intratumor heterogeneity. First, hypothesizing that differences in the number and strength of driver genes contribute to different degrees of intratumor heterogeneity among cancer types, we focused on parameters *d* and *f*, which control driver gene size and strength, respectively. A heat map of *ϵ* in Figure 2A (hereafter, called a *d-f* heat map) showed that a multiple number of driver genes of moderate strength (e.g., 8 ≤ *d* ≤ 10, 0.4 ≤ *f* ≤ 0.5) lead to high intratumor heterogeneity. On the other hand, when cells harbor any strong driver gene, i.e., *f* is high, intratumor heterogeneity is low for any *d*. These observations can be interpreted by focusing on different modes of selective sweeps, which are phenomenons of positive natural selection driving alleles to fixation. If a cell has a strong driver gene, a clone that acquires the driver mutation rapidly dominates in the population, yielding a clonal tumor (Figure 3). In contrast, if cells harbor multiple driver genes of moderate strength, cells gradually increase their growth rate, which provides cells with chances to acquire different combinations of driver mutations and accompanying passenger mutations (Figure 4). These two different modes of selective sweeps are hereafter referred to as the *clonal* and *subclonal selective sweep,* respectively, and the heterogeneity generated by subclonal selective sweep is referred to as *selective sweep-derived heterogeneity.* Note that when cells with small *d* or *f* cannot accumulate driver mutations necessary for clonal expansion, the ‘almost normal’ cells proliferate until population size reaches at *c_max_*. This case is marked by a lack of founder mutations, low population fitness and longer growth time, as shown by *d-f* heat maps of *ρ, λ*, and *τ* (Figure 2B).

**Figure 2:**
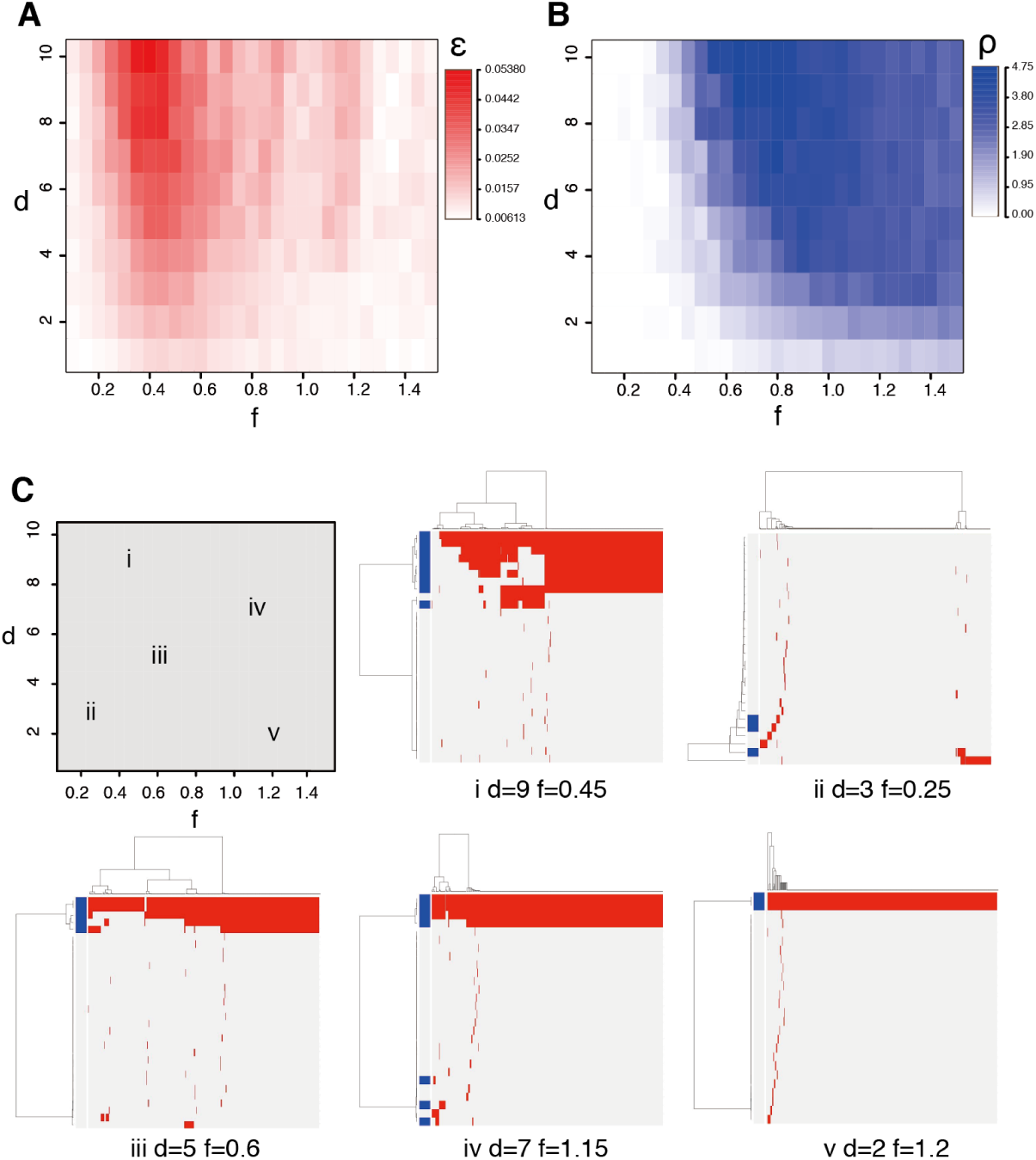
Statistics and mutation profiles. (A) and (B) Population entropy *ϵ* and the number of founder mutations *ρ* were measured for different combinations of *d* and *f* and shown as *d-f* heat maps (C) Representative mutation profiles from simulations with indicated combinations of *d* and *f*. Rows and columns of the mutation profile matrices index genes and cells, respectively, and blue bars denote driver genes.

**Figure 3:**
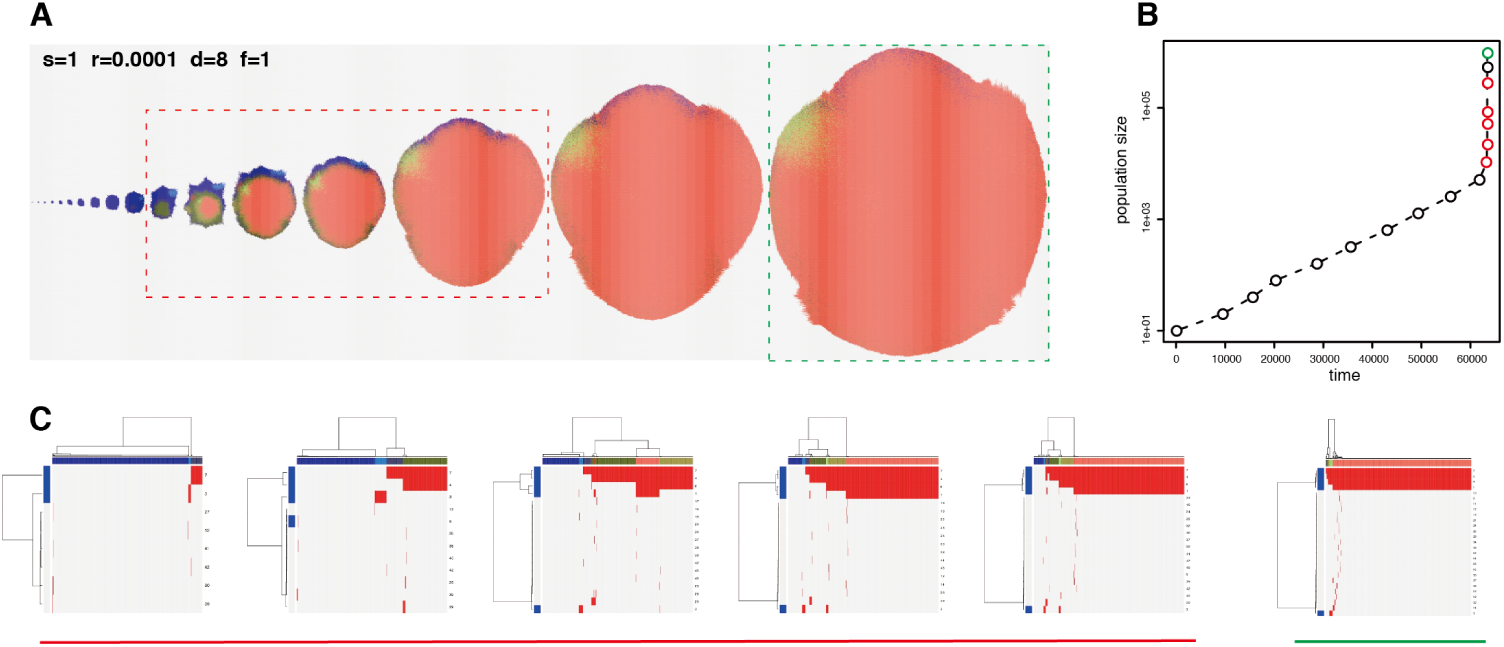
Clonal selective sweep. (A) Snap shots of growing tumors in a simulation with indicated parameter sets. Differently colored cell populations represent each clone. (B) A growth curve of the simulated tumor. The snap shots were obtained at each plotted point. (C) Mutation profiles of the simulated tumor during growth (indicated by red dashed rectangles, plotted points and under lines) and the end point (indicated by green dashed rectangles, plotted points and under lines). Colored bars on the columns represent each clone.

**Figure 4:**
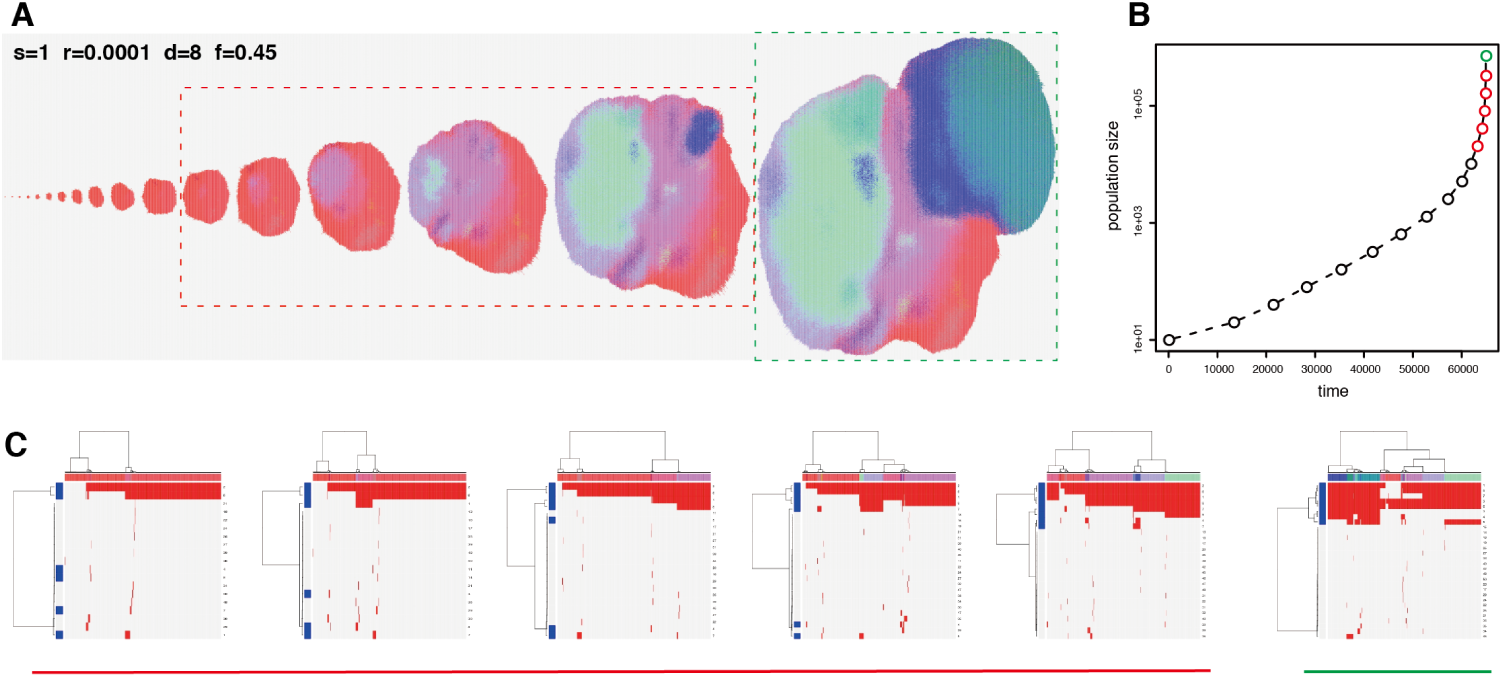
Subclonal selective sweep. shown as in Figure 3.

### Effects of mutation rate

It is also natural to assume a high mutation rate as a cause of intratumor heterogeneity. In addition to a low mutation rate *r* = 0.0001, which is a setting used for the above-mentioned analysis, we tested increased mutation rates, *r* = 0.001 and *r* = 0.01; as expected, we found that an increase in *r* leads to a marked increase in *ϵ* on the average of the *d-f* space (Figure 5A). Additionally, we found that dependence of *ϵ* on *d* and *f* changes with increasing *r*; the parameter dependency observed when *r* = 0.0001 appears to vanish when *r* = 0.001. On the other hand, when *r* is increased to 0.01, a different parameter dependency appears, where *ϵ* is relatively high in areas with low *d* or low *f*. Moreover, cluster heat maps of mutation profile matrices demonstrated that cell-wise dendrograms tended to show fractal-like patterns harboring self-similarity, as *r* increased (Figure 6C). In fact, we calculated the rate of cell-wise dendrograms having such self-similarity (hereafter represented by *θ*) for each parameter setting, and found that *θ* is almost 1 when *r* = 0.01 (Figure 5B). We termed intratumor heterogeneity observed in such cases *fractal heterogeneity* [27]. Close inspection of mutation profile matrix heat maps suggested that the fractal heterogeneity was caused by neutral evolution [24]; that is, numerous neutral mutations that do not affect growth rate were produced by hypermutation and diverse subclones harboring different neutral mutations underwent genetic drift (Figure 7). Note that, depending on *d* and *f*, we observed substantial variations of fractal heterogeneity: e.g., some have large blocks of neutral mutations in the mutation profile (e.g, Figure 6C iii d=6, f=0.6), but others do not (e.g, Figure 6C v d=2, f=1.3). The number of funder mutations is also dependent on the parameters as shown by the *d-f* heat maps of *ρ* (Figure 6B and the supplementary web site).

**Figure 5:**
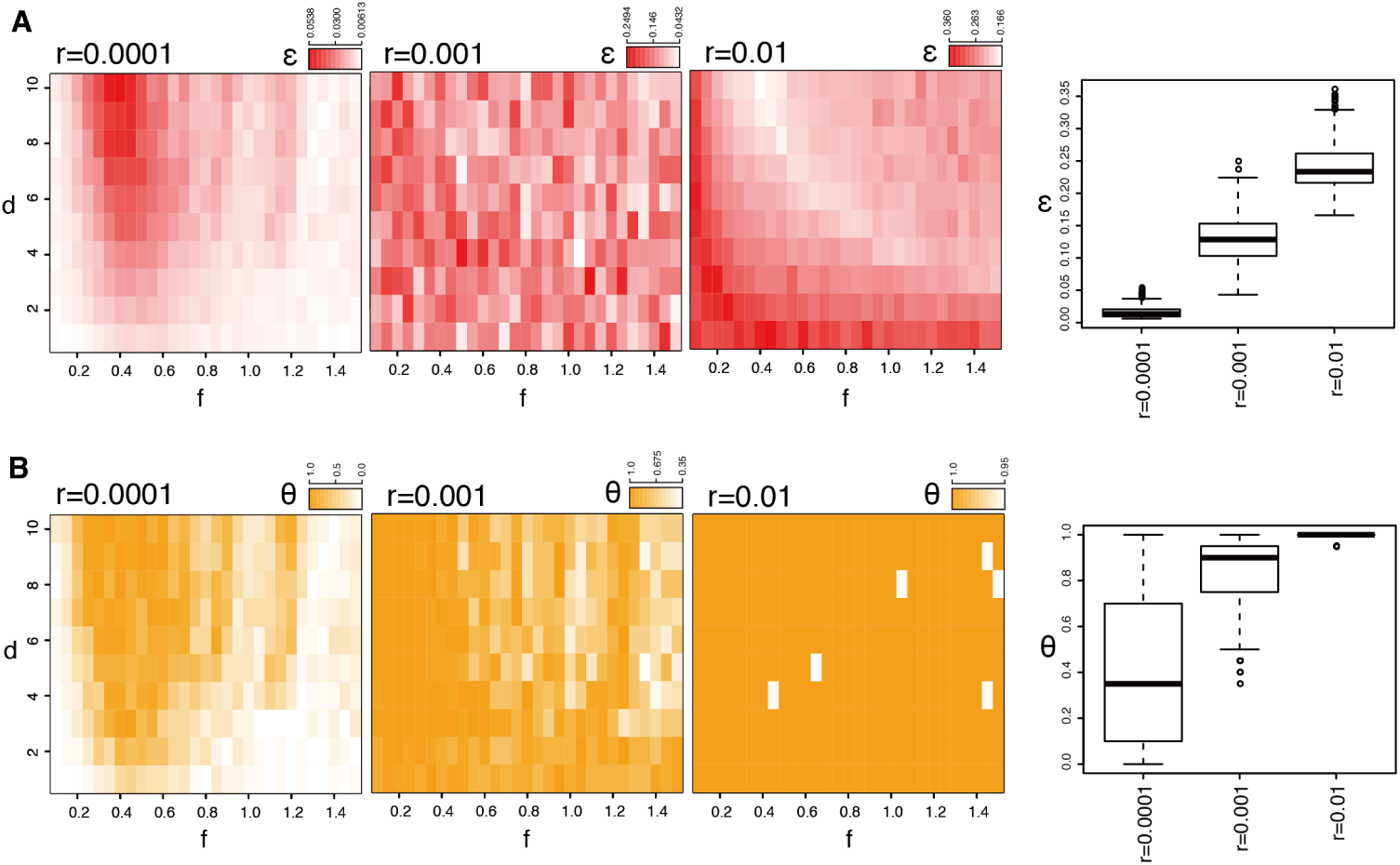
Transitions of heterogeneity caused by an increase of mutation rate. (A) Three left *d-f* heat maps shows Population entropy *ϵ* from simulations with three different values of *r*. Note that scales are different for each heat map. The distributions of the values in each *d-f* heat map is shown in the right box plot. (B) self similarity rate of cell-wise dendrograms *θ* is shown as in A.

**Figure 6:**
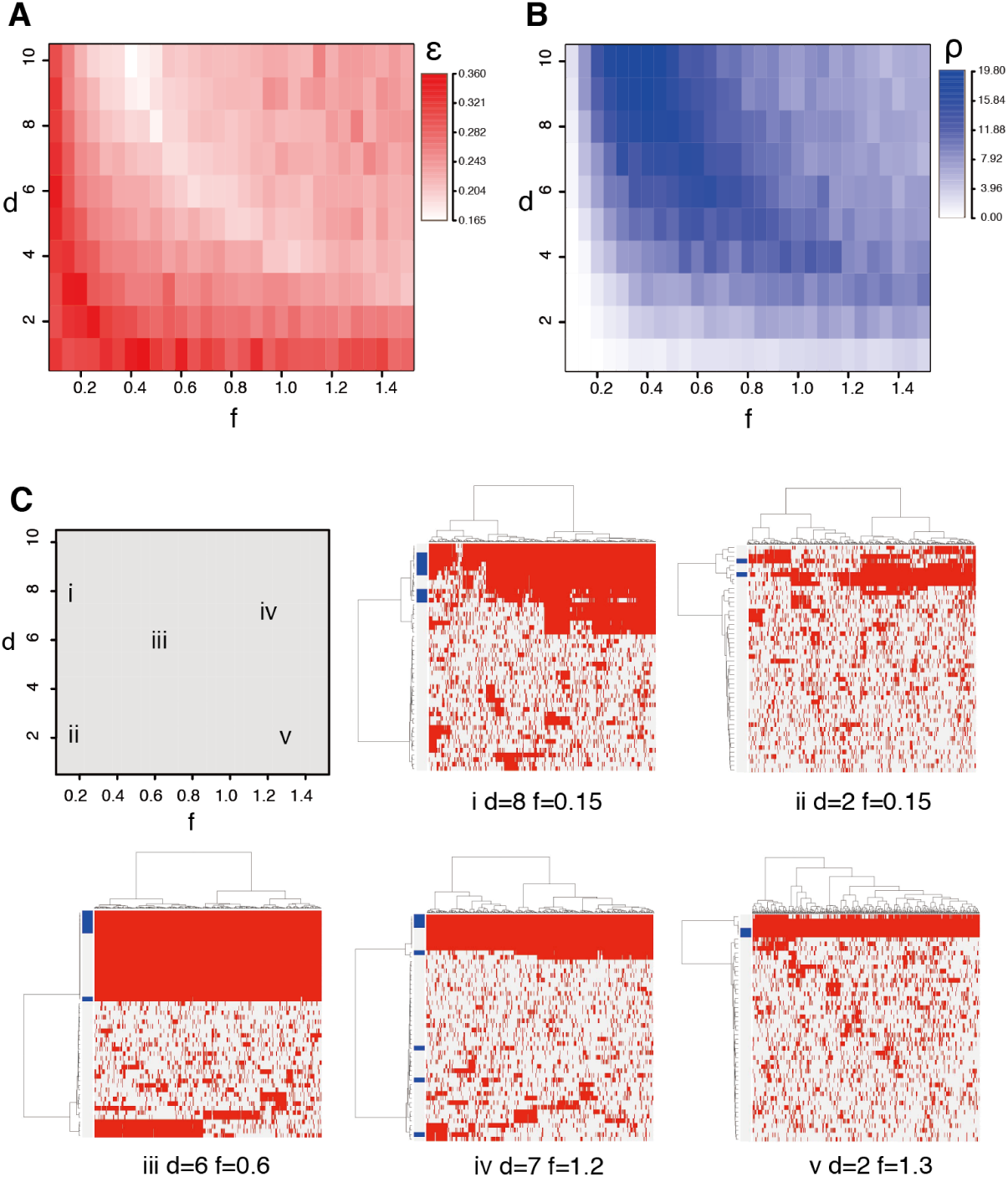
Statistics and mutation profiles from simulation with a high mutation rate. Results from rom simulation with *r* = 0.01 are shown as in Figure 2.

**Figure 7:**
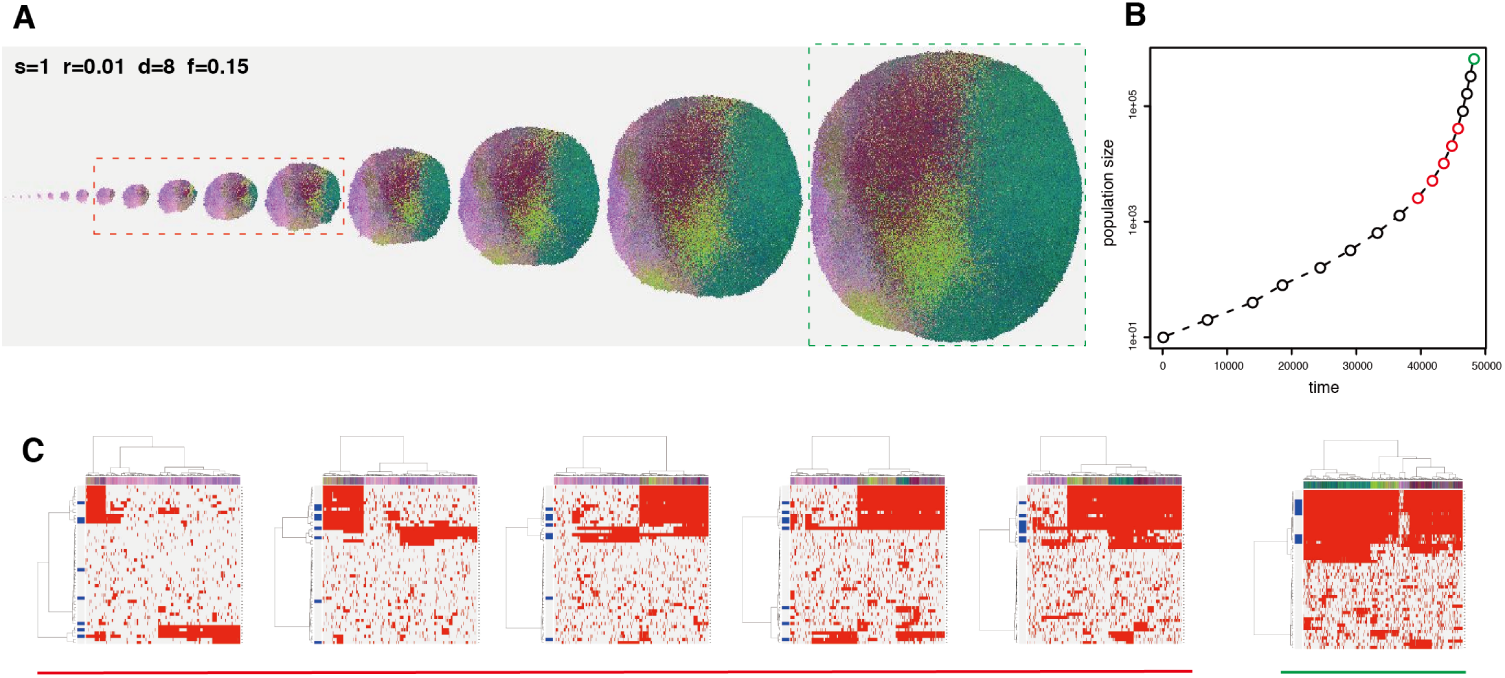
Neutral evolution generating fractal heterogeneity. shown as in Figure 3.

### Effects of a stem cell hierarchy

Over the last decade, many studies accumulated evidence that cancer cells have a stem cell hierarchy, similarly to normal cells [30]. By definition, the stem cell hierarchy generates intratumor heterogeneity in the functional and phenotypic contexts, while this study focused on intratumor heterogeneity in the genomic context. The cancer stem cell concept is also well studied by mathematical modeling [35, 1, 49, 47, 26]. Among them, several studies have suggested a stem cell hierarchy as a mechanism generating genomic intratumor heterogeneity [37, 40]. Inspired by these studies, we extended the BEP model so that it harbors a simple stem cell hierarchy, and examined its effect on genomic intratumor heterogeneity (Figure 8). The extended BEP model assumes two types of cells: stem cells and differentiated cells. Stem cells divide symmetrically and asymmetrically, and the rate of symmetric division is specified by parameter *s*. Differentiated cells, which are produced by asymmetric division, have a shorter life span than stem cells. Namely, the death probability of differentiated cells *q_d_* is larger than that of stem cells *q*_0_. Note that when *s* =1, the extended model does not have a stem cell hierarchy and becomes identical to the model employed so far.

**Figure 8:**
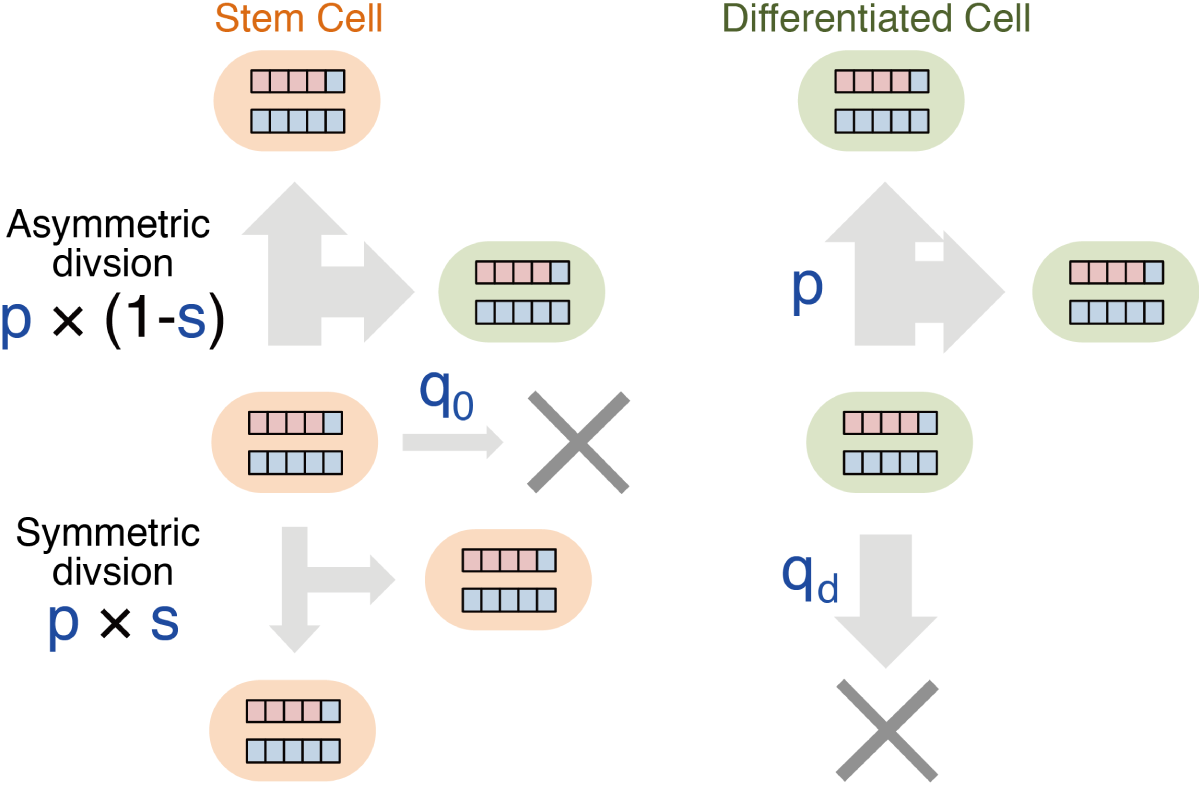
A schema of the BEP model with a stem cell hierarchy. A stem cell divides symmetrically or asymmetrically, and the rate of symmetric division is specified by parameter *s*. The asymmetric division produces a differentiated cell, whose death probability *q_d_* is larger than that of stem cells *q*_0_. The model with *s* =1 is identical to the model without a stem cell hierarchy shown in Figure 1.

We evaluated how a decrease in s (i.e., enhancement of the stem cell hierarchy) affects *ϵ* and *θ*, as evaluated for mutation rate *r* (Figure 9). We found that the dependency of *ϵ* on *d* and *f* varies with decreasing *s*. With *s* = 0.01, A fraction of the *d-f* parameter space starts to increase *ϵ* and, with *s* = 0.01, the parameter space is divided into two distinct areas with high and low *ϵ* (Figure 10A). In the large area associated with high *d* and high *f*, *ϵ* scored low, which was caused by clonal or subclonal selective sweeps. On the other hand, the smaller area associated with low *d* and low *f*, *ϵ* scored distinctively high, and the *d-f* heat map of *θ* suggests that this was derived from fractal heterogeneity. Mutation profiles also show that the parameter area generated heterogenous tumors similar to those generated by the models with high mutation rates (Figure 10C). However, if *d* or *f* is small, the cell populations could not attain high growth rate enough to reach the size limitation *c_max_* before simulation stopped. Collectively, our data demonstrated that, dependent on *d* and *f*, the model with a stem cell hierarchy could generate fractal heterogeneity.

**Figure 9:**
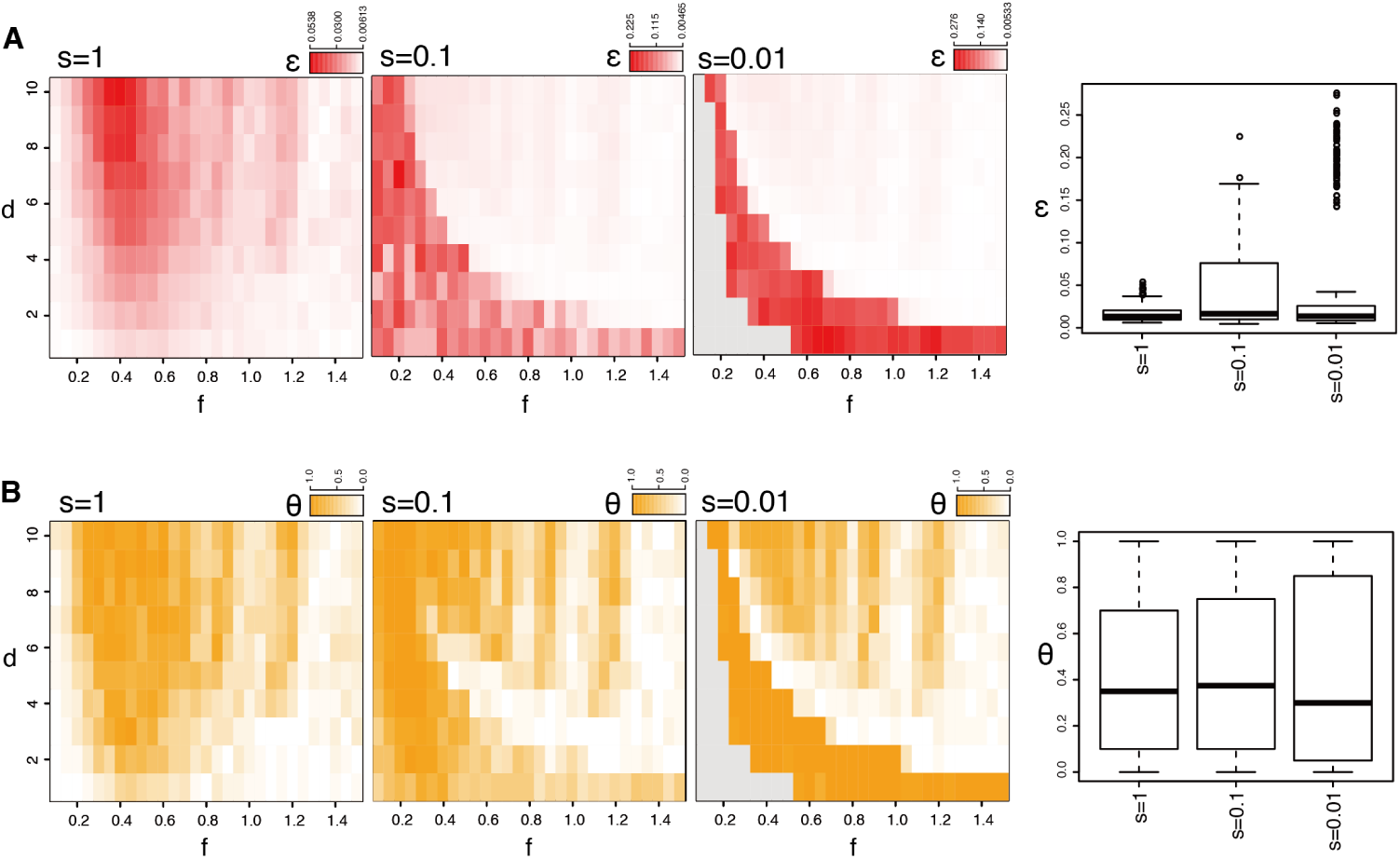
Transitions of heterogeneity caused by enhancing a stem cell hierarchy. Transitions of Population entropy e and self similarity rate *θ* in response to changes of the rate symmetric division rate s are shown as in Figure 5. Gray areas in *d-f* heat maps represent cases where tumors grow slowly and simulations stopped before population size reaches *c_max_*.

**Figure 10:**
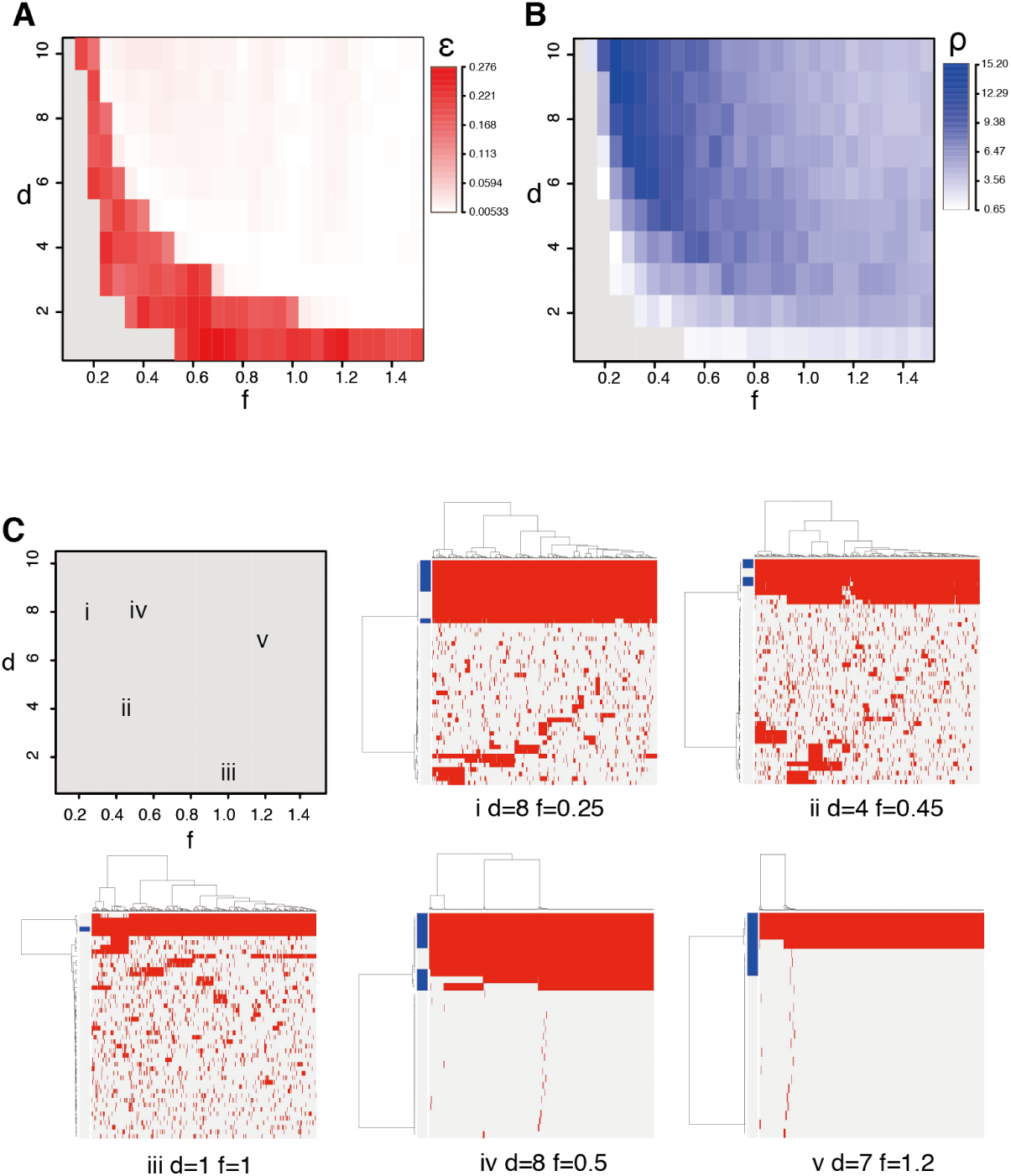
Statistics and mutation profiles from simulation with a stem cell hierarchy. Results from simulation with *s* = 0.01 are shown as in Figure 2. Gray areas in *d-f* heat maps indicate that simulations stopped before population size reaches *c_max_*.

## Disscussion

Here, by simulating branched cancer evoution, we identified three important factors that could be critical for generation of intratumor heterogeneity: 1) the number and strength of driver genes; 2) mutation rate; and 3) a stem cell hierarchy. First, our BEP simulation demonstrated that the number and strength of driver genes are important factors controlling two modes of selective sweeps: clonal and subclonal selective sweeps. If a cell obtains any strong driver mutations, a clone derived from the cell rapidly becomes dominant in the population leading to a clonal tumor. On the other hand, if a cell harbors multiple driver genes of moderate strength, the cell gradually accumulates multiple combinations of mutations and generates selective sweep-derived heterogeneity. Our BEP simulation also showed that a high mutation rate could contribute to extremely high intratumor heterogeneity characterized by fractal patterns of cluster dendrograms; therefore, we termed it fractal heterogeneity. Finally, we extended the BEP model to find that fractal heterogeneity can also be generated by a stem cell hierarchy.

It is a well established fact that cancer genomes accumulate multiple driver mutations during carcinogenesis [19, 51]. The necessity of multiple driver mutations for carcinogenesis could delay development of cancer; it acts as a tumor suppressive mechanism in multicellular organisms. Intriguingly, it has been reported that a stem cell hierarchy could also play an important role in suppressing the development of cancer [31]. Simulations assuming a stem cell hierarchy also required more time steps than those not assuming a stem cell hierarchy, especially for parameter settings generating heterogeneous tumors (see the supplementary web site). Taken together with these facts, our observation suggests that, ironically, intratumor heterogeneity is an inevitable consequence of the implementation of these tumor suppressive mechanisms. In other words, there is a trade-off relationship between intratumor heterogeneity and these tumor suppressive mechanisms.

Acceleration of the mutation rate is a hallmark of cancer [19, 51]; as intuitively expected, our BEP simulation showed that the mutation rate is strongly associated with intratumor heterogeneity. Notably, we found that an increase in the mutation rate leads to extremely high intratumor heterogeneity, which demonstrates a fractal structure. Fractal heterogeneity is generated by neutral evolution; numerous subclones are generated by accumulating neutral mutations and undergo genetic drift. This is in contrast to selective sweep-derived heterogeneity, where a limited number of subclones are generated by positive natural selection. We found that a stem cell hierarchy can similarly produce the fractal heterogeneity. Although further studies are needed to understand why a stem cell hierarchy can generate fractal heterogeneity, one possibility is that the decrease in population size accelerates genetic drift, which is as the founder effect [6, 40].

Our analysis of parameter dependence unveiled transitions between different phases of cancer evolutionary dynamics. For example, an increase of *r* drastically rises *ϵ* and *θ*, which represents a transition from clonal/subclonal selective sweep to neutral evolution generating fractal heterogeneity (Figure 5). A *d-f* heat map of *ϵ* with *s* = 0.01 also shows a clear boundary between the two distinct phases (Figure 10A). Intriguingly, it has been reported that complex systems often undergo such drastic phase transitions as observed in our data. Langton found that, while changing a system parameter, cellular automata interchange between highly ordered and highly disordered dynamics, and termed their boundary the edge of chaos [25]. Kauffman studied mathematical models of evolving systems and proposed that the rate of evolution is maximized near the edge of chaos [23]. In the BEP model, the clear phase boundary in the *d-f* heat map of *ϵ* with *s* = 0.01 (Figure 10A) could be compared to the edge of chaos. Taken together, it is tempting to hypothesize that cancer is also evolving at the edge of chaos, which underlies its extraordinary evolvability.

As stated in the introduction, recent multiregional genomic studies have revealed highly branched evolution in a number of solid tumors; renal clear cell carcinoma and low-grade glioma tend to have subclonal driver mutations, which are often located in different positions of known driver genes [16, 15, 44]. Although this observation may support selective sweep-derived heterogeneity, most of the terminal branches in the evolutionary trees lack clear subclonal driver mutations. It has been also reported that pancreatic tumors harbor established driver mutations only as founder mutations, although many progressor mutations accumulate before metastasis occurs [50]. Recently, multiregional sequencing of non-small cell lung cancer is reported; lung cancer appeared to harbor founder mutations only in founder mutation [52, 8]. Similarly, we performed multiregional sequencing of colorectal cancers to find that colorectal cancer evolution demonstrates highly branched patterns without clear subclonal driver mutations. Moreover, when a parameter set leading to neutral evolution is used, in silico multiregional sequencing of a simulated tumor from the BEP model well reproduced the experimental result (Uchi *et al.* in submission). Another recent multiregional study of colorectal cancer also showed an absence of selective sweeps [39]. From these observations, we hypothesize that branched cancer evolution is mainly drived by neutral evolution and intratumor heterogeneity is essentially fractal. Fractal heterogeneity also appeared to be consistent with emerging evidence that resistance to some targeted cancer drugs may result from the outgrowth of preexisting low frequency cancer cell populations [9]. If a tumor has fractal heterogeneity, numerous subclones with different neutral mutations would expand a repertoire of potentially resistant clones in the presence of therapeutic selection. This point is clinically important and should be addressed in detail in future studies.

In this study, we employed a cellular automaton model to simulate branched cancer evolution. Although there have exist many excellent cellular automaton models models that reproduce various sides of humor biology like microenvironmental interaction and cellular movement [22, 21, 46, 14, 2, 45], we stuck to make the BEP model as simple as possible, aiming to explore principles behind branched evolution. The simplification of the model is necessary to suppress the curse of dimensionality in the parameter dependence analysis on our supercomputer, which successfully provided us insights into the principles. However, our analysis is far from covering the whole of the parameter space; more fine-grained parameter analyses are needed to obtain a complete phase diagram of the evolutionary dynamics. Another limitation of this study should also be noted: we set some of the parameter values to different orders of magnitude from realistic values. For example, we set genome size *n* = 50, but the human genome has 2 × 10^4^ genes and 3 × 10^9^ bases. Corresponding to the smaller number of genes, we also use high mutation rate rate (*r* = 10^−4^−10^−2^), compared to that assumed in previous works [3, 53, 5]. We simulated tumor growth until the population size reached *c_max_* = 10^6^; however, the actual tumor size is estimated from 10^9^ to 10^10^. Memory limitation forced us to use these unrealistic parameter settings: because even a model with *n* = 50 can produce the virtually infinite number of the branched states, even keeping the branched stets of millions of cell needs several gigabytes of memory in our current implementation. To overcome these problems, better implementation of the BEP model or analytical solution of the BEP model are awaited.

Moreover, although we believe that our simple model catches the essence of branched cancer evolution, many points should be improved to to obtain a more realistic model. For example, although we assumed that each driver mutations has equal and independent effects on growth rate, the effects should be different genes, epistatic gene interactions should exist, and some of the driver mutations should affect death rate. It is also probable that the parameters could change in the spatial and temporal scales of tumor evolution: the mutation rate would be elevated after caretaker genes are mutated and genomic instability is incurred. Assuming these conditions, we propose that a tumor first undergoes clonal/subclonal selective sweep, and later expands heterogeneously by neutral evolution. There would also exist interactions between cells and microenvironments; a mutation conferring a growth advantage in a specific microenvironment could be a subclonal driver mutation, further enhancing intratumor heterogeneity. Competition and interaction between cells should be considered and a real tumor grows in a three dimensional space. we will address these extension of the BEP model as a future work.

In conclusion, this study introduced the BEP model, a novel mathematical model of cancer evolution. Moreover, employing our supercomputer, we performed the extensive parameter dependance analysis of the BEP model, which brought a number of intriguing insights into branched cancer evolution. Among these, the hypothesis that cancer has fractal heterogeneity could be the most important key for understanding cancer evolvability; this awaits experimental validation. Combined with recent advance in experimental technologies, such as single cell sequencing [48], our BEP simulation will be a insightful tool for understanding heterogenous cancer evolution and the resulting malignancy.

## Method

### Simulating cancer evolution

To simulate branched cancer evolution, we employed a cellular automaton model, assuming that a cell in the tumor acts as a cellular automaton. A cell has a genome containing *n* genes, each of which is represented as a binary value, 0 (wild) or 1 (mutated). Namely, the genome is represented as a binary vector **g** of length *n*. In a unit time step, each cell in the simulated tumor dies with a probability *q*. If the cell does not die, the cell then divides with a probability *p*. Before the cell division, we mutate the genome vector *g*: each of 0 elements of *g* is set to 1 with a probability *r*. The first *d* genes in **g** are assumed as driver genes, whose mutations accelerate the division speed. A normal cell without any mutations has a division probability *p*_0_, and acquisition of one driver mutation increases *p* by 10*^f^*-fold in the next time step; i.e., *p* = *p*_0_ *·* 10*^fk^*, where 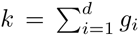 the number of mutated driver genes. The death probability is fixed as q = *q*_0_. Let *c* and *t* denote the size of the simulated cell population and the number of the time steps, respectively. We started a simulation with *c*_0_ normal cells and repeated the unit time step while population size *c* ≤ *c_max_* and time step *t* ≤ *t_max_.* A flowchart of the simulation is shown as Figure 11A.

**Figure 11:**
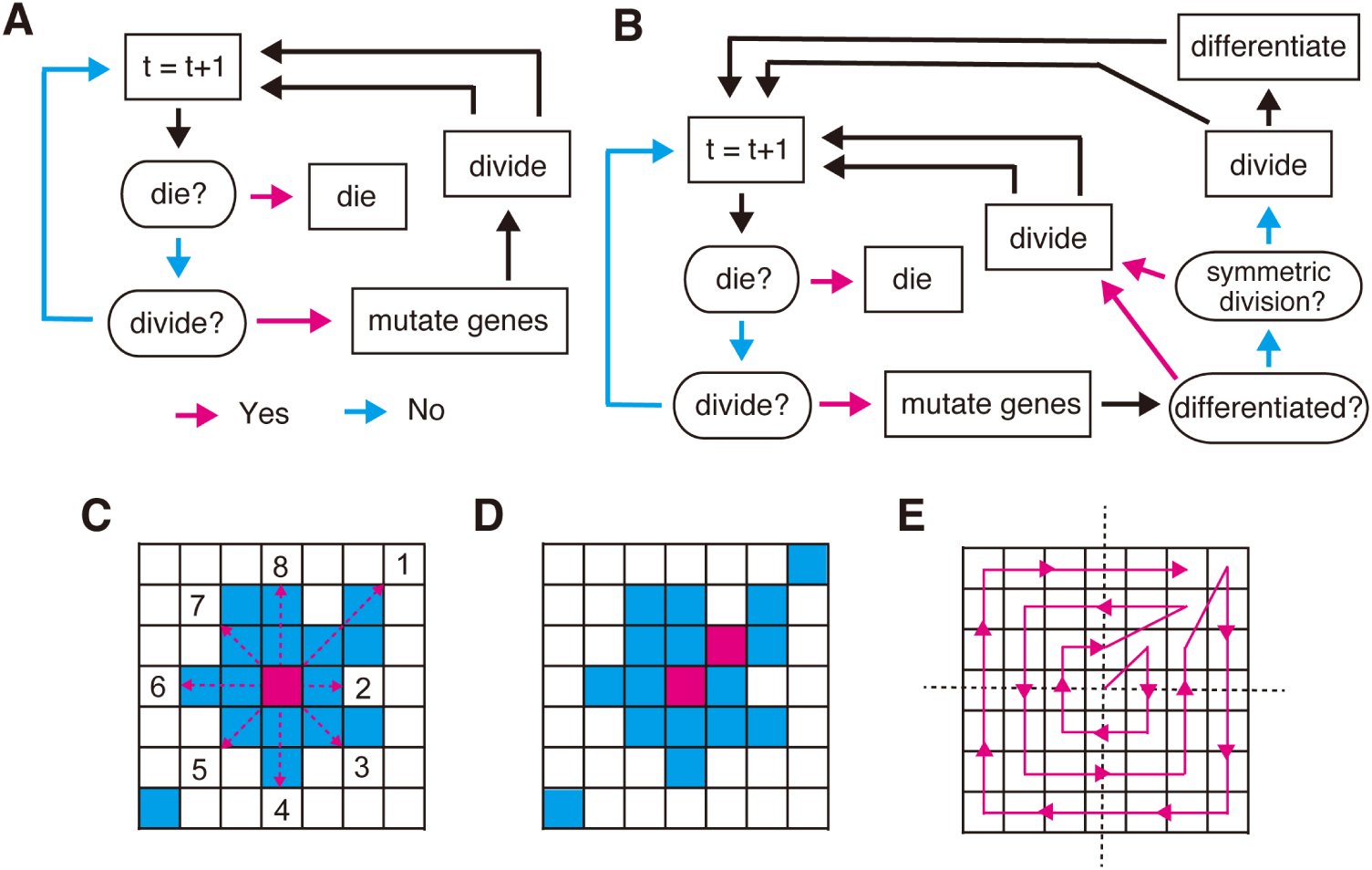
Illustration of our simulation method. (A,B) Flowcharts of our simulation without (A) and with a stem cell hierarchy (B). In each time step, first, cell dies with a probability *q* and, if the cell survives, the cell divides with a probability *p*. Before the cell division, the genome is subject to mutagenesis. In simulation with a stem cell hierarchy, a differentiated cell to be divided always undergoes symmetric division. a stem cell to be divided undergoes symmetric division with a probability *s*. Otherwise, it undergoes asymmetric division, which generates a stem cell and a differentiated cell. (C, D, E) Division operation. We randomly select one point from in the eight neighbor points, if empty points exist. Otherwise, we create an empty point neighboring the targeted point (a red point in C) by: 1) for each of the eight directions (red dotted arrows in C), obtaining *l_i_* (1 ≤ *i* ≤ 8), the number of the consecutive occupied points that range from each neighboring points to immediately before the nearest empty cell (numbered points in C), and 2) selecting either of the 8 directions with a probability proportional with 1/*l_i_*. In the case shown in C, *l*_1_ = 2, *l*_2_ = 1, *l*_3_ = 1,... The consecutive occupied points in the selected direction are then shifted by one point so that an empty neighboring point appears. A case where the first direction is selected in C is shown in D. This operation is applied to each cell to be divided along an outward spiral starting from the center, whose direction is randomly flipped in each round. E shows an example of such a spiral.

Moreover, we incorporated the cancer stem cell concept into the BEP model. We assumed two different types of cells: stem cells and differentiated cells. we assumed that a differentiated cell has a shorter life span, compared to a stem cell: the death probability of stem cells is *q*_0_ while that of differentiated cells is *q_d_ > q*_0_. A stem cell divides asymmetrically to produce one stem cell and one differentiated cell, or otherwise symmetrically to produce two stem cells. Namely, a stem cell to be divided divides symmetrically and asymmetrically with probabilities s and (1 − *s*), respectively. A differentiated cell divides only symmetrically to produce two differentiated cells. Note that when *s* = 1, the model becomes identical to the model without a stem cell hierarchy. A flowchart of the simulation with stem a cell hierarchy is shown as Figure 11B.

### Visualizing simulated tumors

We designed a simulated tumor to grow in a two-dimensional square lattice where each cell occupies one lattice point. In the beginning, *c*_0_ cells are initialized as close as possible to the center of the lattice. when a cell die, the occupied point is cleared and becomes empty. When a cell divides, we place the daughter cell in the neighborhood of the parent cell, assuming the Moore neighborhood (i.e., eight points surrounding a central point). If empty neighbor points exist, we randomly select one point from them. Otherwise, we create an empty point in either of the eight neighboring points by the following procedure. First, for each of the eight directions, we count the number of the consecutive occupied points that range from each neighboring points to immediately before the nearest empty cell as indicated in Figure 11C. Next, either of the eight directions is selected with a probability proportional with 1/*l_i_*, where *l_i_*(1 ≤ *i ≤* 8) is the count of the consecutive occupied points for each direction. The consecutive occupied points in the selected direction are then shifted by one point so that an empty neighboring point appears as shown in Figure 11D. Note that simulation results are dependent on the order of the division operation in the two-dimensional square lattice. We first marked cells to be divided, and applied the division operation to the marked cells along an outward spiral starting from the center. In each round on the spiral, the direction was randomly flipped in order to keep spatial symmetry. An example of such spirals was shown in Figure 11E.

From a simulated tumor, we randomly sampled m cells to obtain an n × m mutation profile matrix **M**, each of whose *m* columns is the *n*-dimensional genome vector **g** for each cell. We applied principle component analysis to **M** and obtained the first, second and third loading vectors. By multiplying these loading vectors, *n*-dimensional genome vectors were reduced into threedimensional vectors. RGB colors used for sample labels are then prepared by mixing red, green and blue proportionally to the three vector elements. Each cell on the two-dimensional square lattice was colored with the color corresponding to its genotype. **M** was visualized using a cluster heat map, to which we appended a colored bar representing the genotype of each cell.

### Calculating statistics of simulation results

To quantify intratumor heterogeneity, we defined a population entropy statistic, *ϵ*, based on our previously reported approach for measuring information content in a matrix [33]. From **M**, we calculated similarities among m cells and obtained a similarity matrix **S**. Similarity *S_ij_* between two cells ***g****^i^* and ***g****^j^* is calculated as 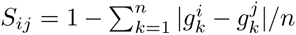 We applied singular value decomposition to **S** and obtained a singular value vector **s**. We then defined *ϵ* by 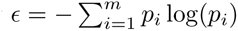, where 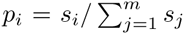.*ϵ* is 0 when the population is composed of one clone, but increase with genetic variation of the population.

Similarly, a number of statistics were obtained to evaluate simulation results. From the mutation profile matrix **M**, we obtained the average mutation count as follows: 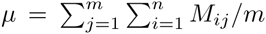. Assuming mutations occurring in more than 95 percent of a population as founder mutations, we also calculated the average founder mutation count: 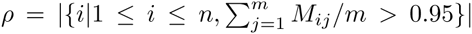. Population fitness was measured by: 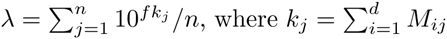, and the number of time steps required for the population size *c* to reach *c_max_* was recorded as *τ*.

### Testing self-similarity of mutation profiles

We performed hierarchical clustering of randomly sampled *m*′ cells by applying Ward’s method to a distance matrix 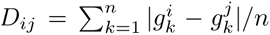 A cluster dendrogram is then obtained and examined for self-similarity. Our approach is based on a ‘box-counting’ method for measuring fractal dimensions [38]. Given *a*, which is less than the height of the dendrogram, we can obtain *b* clusters by cutting the dendrogram at the height *a*. If the dendrogram exhibits self similarity, the following relationship holds: *b* ∝ *a*^−6^, where *δ* is a fractal dimension. To evaluate whether the dendrogram has this relationship, we obtained *a* from 1000 points with equal intervals between the minimum and maximum height to calculate *b* for each *a*. We then performed linear regression between log(*a*) and log(*b*), and assumed that the dendrogram has self-similarity if *R*^2^ > 0.95

### Analyzing parameter dependence

To examine the parameter dependance of each statistic, we varied *d, f*, *r* and *s* while others were fixed, as summarized in Table 1. We prepared 10 integers from 1 to 10 as *d*; 29 numbers from 0.1 to 1.5 incremented by 0.05 as *f*; 1, 0.1 and 0.01 as *s*; and 0.01, 0.001 and 0.0001 as *r*, and take every combinations of the parameter values as done in grid search. This leads to 10 × 29 × 3 × 3 = 2610 parameter settings, for each of which simulation was repeated 20 times. 2610 × 20 = 52200 simulations were performed parallelly on our supercomputer system [32]. For each parameter setting, we obtained the averaged statistics of the 20 simulations. The proportion of simulated tumors whose cell-wise dendrograms have self-similarity was also obtained as a statistic, *θ*. The obtained statistics are summarized in Table 2. The results of the parameter dependence analysis were subject to interactive visualization using the D3 JavaScript library (http://d3js.org/), which can be assessed at the supplementary website: http://bep.hgc.jp/.

**Table 1:**
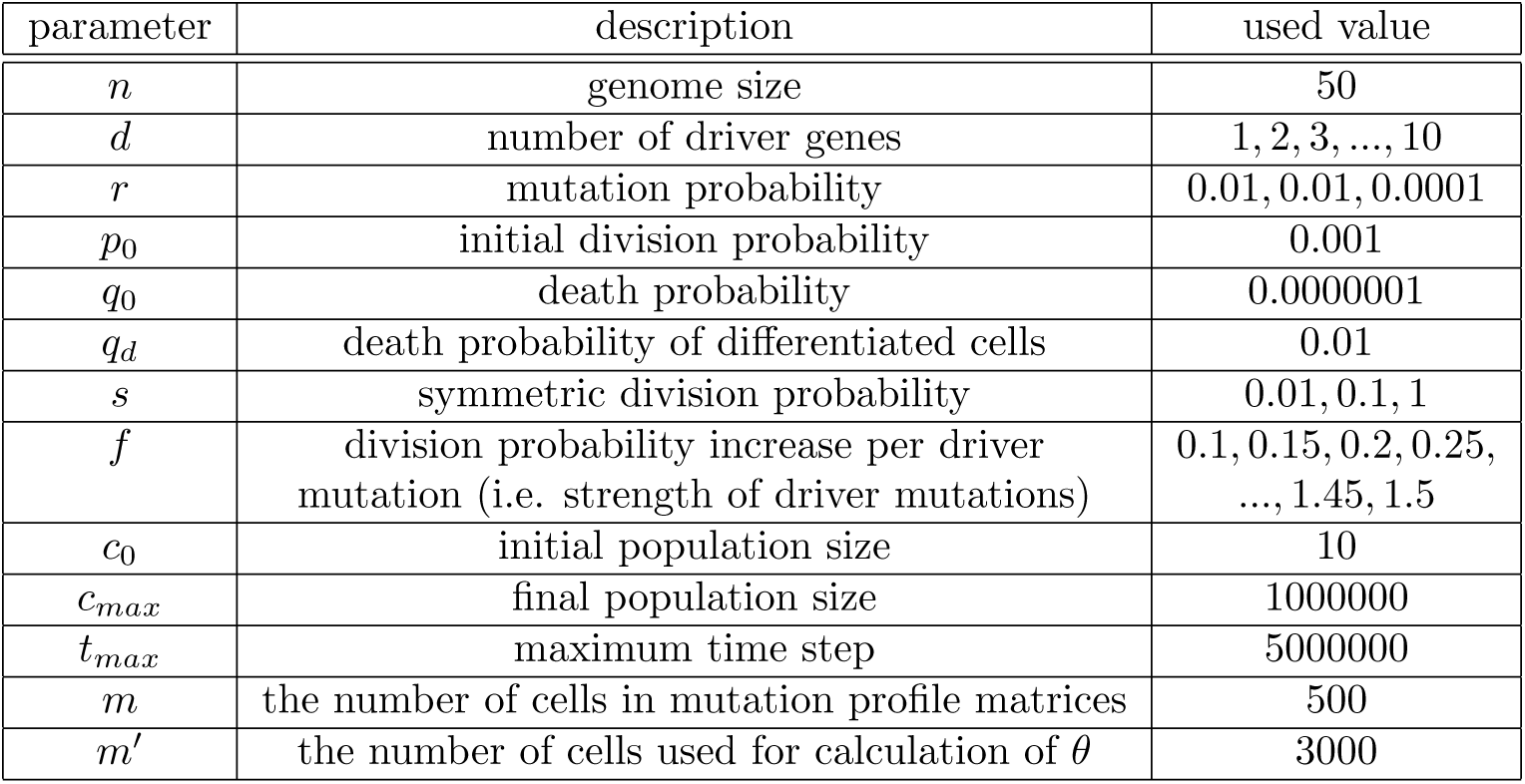
Parameter values used in this study

| parameter | description | used value |
| --- | --- | --- |
| $n$ | genome size | 50 |
| $d$ | number of driver genes | 1, 2, 3, ..., 10 |
| $r$ | mutation probability | 0.01, 0.01, 0.0001 |
| $p_0$ | initial division probability | 0.001 |
| $q_0$ | death probability | 0.0000001 |
| $q_d$ | death probability of differentiated cells | 0.01 |
| $s$ | symmetric division probability | 0.01, 0.1, 1 |
| $f$ | division probability increase per driver mutation (i.e. strength of driver mutations) | 0.1, 0.15, 0.2, 0.25, ..., 1.45, 1.5 |
| $c_0$ | initial population size | 10 |
| $c_{max}$ | final population size | 1000000 |
| $t_{max}$ | maximum time step | 5000000 |
| $m$ | the number of cells in mutation profile matrices | 500 |
| $m'$ | the number of cells used for calculation of $\theta$ | 3000 |

**Table 2:** Statistics of simulation results

| statistics | description |
| --- | --- |
| $\epsilon$ | population entropy |
| $\mu$ | average count of mutations per cell |
| $\rho$ | count of founder mutations |
| $\lambda$ | population fitness |
| $\tau$ | number of time step needed for tumor growth |
| $\theta$ | self similarity rate of cell-wise dendrograms |

## Acknowledgments

We thank to Drs Gouhei Tanaka, Hiroshi Haeno, and Kimiyo Yamamoto for helpful discussion. This research used computational resources of the K computer provided by the RIKEN Advanced Institute for Computational Science through the HPCI System Research project (Project ID: hp140230). Computation time was also provided by the Super Computer System, Human Genome Center, Institute of Medical Science, University of Tokyo.

## References

[1] Z Agur, OU Kirnasovsky, G Vasserman, L Tencer-Hershkowicz, Y Kogan, H Harrison, R Lamb, and RB Clarke. Dickkopf1 regulates fate decision and drives breast cancer stem cells to differentiation: an experimentally supported mathematical model. PLoS One, 6:e24225, 2011.

[2] D Basanta, DW Strand, RB Lukner, OE Franco, DE Cliffel, GE Ayala, SW Hayward, and AR Anderson. The role of transforming growth factorbeta-mediated tumor-stroma interactions in prostate cancer progression: an integrative approach. Cancer Res, 69:7111–20, 2009.

[3] N Beerenwinkel, T Antal, D Dingli, A Traulsen, KW Kinzler, VE Velculescu, B Vogelstein, and MA Nowak. Genetic progression and the waiting time to cancer. PLoS Comput Biol, 3:e225, 2007.

[4] N Beerenwinkel, RF Schwarz, M Gerstung, and F Markowetz. Cancer evolution: mathematical models and computational inference. Syst Biol, 64:e1–25, 2015.

[5] I Bozic, T Antal, H Ohtsuki, H Carter, D Kim, S Chen, R Karchin, KW Kinzler, B Vogelstein, and MA Nowak. Accumulation of driver and passenger mutations during tumor progression. Proc Natl Acad Sci U S A, 107:18545–50, 2010.

[6] B Charlesworth. Fundamental concepts in genetics: effective population size and patterns of molecular evolution and variation. Nat Rev Genet, 10:195–205, 2009.

[7] K Danesh, R Durrett, LJ Havrilesky, and E Myers. A branching process model of ovarian cancer. J Theor Biol, 314:10–5, 2012.

[8] Bruin EC de, N McGranahan, R Mitter, M Salm, DC Wedge, L Yates, M Jamal-Hanjani, S Shafi, N Murugaesu, AJ Rowan, E Gronroos, MA Muhammad, S Horswell, M Gerlinger, I Varela, D Jones, J Marshall, T Voet, Loo P Van, DM Rassl, RC Rintoul, SM Janes, SM Lee, M Forster, T Ahmad, D Lawrence, M Falzon, A Capitanio, TT Harkins, CC Lee, W Tom, E Teefe, SC Chen, S Begum, A Rabinowitz, B Phillimore, B Spencer-Dene, G Stamp, Z Szallasi, N Matthews, A Stewart, P Campbell, and C Swanton. Spatial and temporal diversity in genomic instability processes defines lung cancer evolution. Science, 346:251–6, 2014.

[9] LA Jr Diaz, RT Williams, J Wu, I Kinde, JR Hecht, J Berlin, B Allen, I Bozic, JG Reiter, MA Nowak, KW Kinzler, KS Oliner, and B Vogelstein. The molecular evolution of acquired resistance to targeted egfr blockade in colorectal cancers. Nature, 486:537–40, 2012.

[10] R Durrett, J Foo, K Leder, J Mayberry, and F Michor. Evolutionary dynamics of tumor progression with random fitness values. Theor Popul Biol, 78:54–66, 2010.

[11] R Durrett, J Foo, K Leder, J Mayberry, and F Michor. Intratumor heterogeneity in evolutionary models of tumor progression. Genetics, 188:461–77, 2011.

[12] GB Ermentrout and L Edelstein-Keshet. Cellular automata approaches to biological modeling. J Theor Biol, 160:97–133, 1993.

[13] J Gallaher and AR Anderson. Evolution of intratumoral phenotypic heterogeneity: the role of trait inheritance. Interface Focus, 3:20130016, 2013.

[14] P Gerlee and AR Anderson. Evolution of cell motility in an individual-based model of tumour growth. J Theor Biol, 259:67–83, 2009.

[15] M Gerlinger, S Horswell, J Larkin, AJ Rowan, MP Salm, I Varela, R Fisher, N McGranahan, N Matthews, CR Santos, P Martinez, B Phillimore, S Begum, A Rabinowitz, B Spencer-Dene, S Gulati, PA Bates, G Stamp, L Pickering, M Gore, DL Nicol, S Hazell, PA Futreal, A Stewart, and C Swanton. Genomic architecture and evolution of clear cell renal cell carcinomas defined by multiregion sequencing. Nat Genet, 46:225–33, 2014.

[16] M Gerlinger, AJ Rowan, S Horswell, J Larkin, D Endesfelder, E Gronroos, P Martinez, N Matthews, A Stewart, P Tarpey, I Varela, B Phillimore, S Begum, NQ McDonald, A Butler, D Jones, K Raine, C Latimer, CR Santos, M Nohadani, AC Eklund, B Spencer-Dene, G Clark, L Pickering, G Stamp, M Gore, Z Szallasi, J Downward, PA Futreal, and C Swanton. Intratumor heterogeneity and branched evolution revealed by multiregion sequencing. N Engl J Med, 366:883–92, 2012.

[17] I Gonzalez-Garcia, RV Sole, and J Costa. Metapopulation dynamics and spatial heterogeneity in cancer. Proc Natl Acad Sci USA, 99:13085–9, 2002.

[18] H Haeno, Y Iwasa, and F Michor. The evolution of two mutations during clonal expansion. Genetics, 177:2209–21, 2007.

[19] D Hanahan and RA Weinberg. The hallmarks of cancer. Cell, 100:57–70, 2000.

[20] Y Iwasa and F Michor. Evolutionary dynamics of intratumor heterogeneity. PLoS One, 6:e17866, 2011.

[21] Y Jiao and S Torquato. Emergent behaviors from a cellular automaton model for invasive tumor growth in heterogeneous microenvironments. PLoS Comput Biol, 7:e1002314, 2011.

[22] Y Kam, KA Rejniak, and AR Anderson. Cellular modeling of cancer invasion: integration of in silico and in vitro approaches. J Cell Physiol, 227:431–8, 2012.

[23] SA Kauffman. The origins of order: Self-organization and selection in evolution. Oxford university press, 1993.

[24] M Kimura. The neutral theory of molecular evolution. Cambridge University Press, 1984.

[25] CG Langton. Computation at the edge of chaos: phase transitions and emergent computation. Physica D: Nonlinear Phenomena, 42.1:12–37, 1990.

[26] F Li, H Tan, J Singh, J Yang, X Xia, J Bao, J Ma, M Zhan, and ST Wong. A 3d multiscale model of cancer stem cell in tumor development. BMC Syst Biol, 7 Suppl 2:S12, 2013.

[27] BB Mandelbrot. The fractal geometry of nature. Vol. 173. Macmillan, 1983.

[28] EA Martens, R Kostadinov, CC Maley, and O Hallatschek. Spatial structure increases the waiting time for cancer. New J Phys, 13, 2011.

[29] N McGranahan and C Swanton. Biological and therapeutic impact of intratumor heterogeneity in cancer evolution. Cancer Cell, 27:15–26, 2015.

[30] CE Meacham and SJ Morrison. Tumour heterogeneity and cancer cell plasticity. Nature, 501:328–37, 2013.

[31] F Michor, MA Nowak, SA Frank, and Y Iwasa. Stochastic elimination of cancer cells. Proc Biol Sci, 270:2017–24, 2003.

[32] H Miyazaki, Y Kusano, N Shinjou, F Shoji, M Yokokawa, and T Watanabe. Overview of the k computer system. Fujitsu Scientific and Technical Journal, 48:255–265, 2012.

[33] A Niida, S Imoto, R Yamaguchi, M Nagasaki, A Fujita, T Shimamura, and S Miyano. Model-free unsupervised gene set screening based on information enrichment in expression profiles. Bioinformatics, 26:3090–7, 2010.

[34] MA Nowak, F Michor, and Y Iwasa. The linear process of somatic evolution. Proc Natl Acad Sci USA, 100:14966–9, 2003.

[35] J Poleszczuk, P Hahnfeldt, and H Enderling. Evolution and phenotypic selection of cancer stem cells. PLoS Comput Biol, 11:e1004025, 2015.

[36] Datta R S, A Gutteridge, C Swanton, CC Maley, and TA Graham. Modelling the evolution of genetic instability during tumour progression. Evol Appl, 6:20–33, 2013.

[37] RV Sole, C Rodriguez-Caso, TS Deisboeck, and J Saldana. Cancer stem cells as the engine of unstable tumor progression. J Theor Biol, 253:629–37, 2008.

[38] C Song, S Havlin, and HA Makse. Self-similarity of complex networks. Nature, 433:392–5, 2005.

[39] A Sottoriva, H Kang, Z Ma, TA Graham, MP Salomon, J Zhao, P Marjoram, K Siegmund, MF Press, D Shibata, and C Curtis. A big bang model of human colorectal tumor growth. Nat Genet, 47:209–16, 2015.

[40] A Sottoriva, JJ Verhoeff, T Borovski, SK McWeeney, L Naumov, JP Medema, PM Sloot, and L Vermeulen. Cancer stem cell tumor model reveals invasive morphology and increased phenotypical heterogeneity. Cancer Res, 70:46–56, 2010.

[41] A Sottoriva, L Vermeulen, and S Tavare. Modeling evolutionary dynamics of epigenetic mutations in hierarchically organized tumors. PLoS Comput Biol, 7:e1001132, 2011.

[42] SL Spencer, RA Gerety, KJ Pienta, and S Forrest. Modeling somatic evolution in tumorigenesis. PLoS Comput Biol, 2:e108, 2006.

[43] MR Stratton, PJ Campbell, and PA Futreal. The cancer genome. Nature, 458:719–24, 2009.

[44] H Suzuki, K Aoki, K Chiba, Y Sato, Y Shiozawa, Y Shiraishi, T Shimamura, A Niida, K Motomura, F Ohka, T Yamamoto, K Tanahashi, M Ranjit, T Wakabayashi, T Yoshizato, K Kataoka, K Yoshida, Y Nagata, A Sato-Otsubo, H Tanaka, M Sanada, Y Kondo, H Nakamura, M Mizoguchi, T Abe, Y Muragaki, R Watanabe, I Ito, S Miyano, A Natsume, and S Ogawa. Mutational landscape and clonal architecture in grade ii and iii gliomas. Nat Genet, 47:458–68, 2015.

[45] M Tektonidis, H Hatzikirou, A Chauviere, M Simon, K Schaller, and A Deutsch. Identification of intrinsic in vitro cellular mechanisms for glioma invasion. J Theor Biol, 287:131–47, 2011.

[46] S Torquato. Toward an ising model of cancer and beyond. Phys Biol, 8:015017, 2011.

[47] V Vainstein, OU Kirnasovsky, Y Kogan, and Z Agur. Strategies for cancer stem cell elimination: insights from mathematical modeling. J Theor Biol, 298:32–41, 2012.

[48] Y Wang, J Waters, ML Leung, A Unruh, W Roh, X Shi, K Chen, P Scheet, S Vattathil, H Liang, A Multani, H Zhang, R Zhao, F Michor, F Meric-Bernstam, and NE Navin. Clonal evolution in breast cancer revealed by single nucleus genome sequencing. Nature, 512:155–60, 2014.

[49] SL Weekes, B Barker, S Bober, K Cisneros, J Cline, A Thompson, L Hlatky, P Hahnfeldt, and H Enderling. A multicompartment mathematical model of cancer stem cell-driven tumor growth dynamics. Bull Math Biol, 76:1762–82, 2014.

[50] S Yachida, S Jones, I Bozic, T Antal, R Leary, B Fu, M Kamiyama, RH Hruban, JR Eshleman, MA Nowak, VE Velculescu, KW Kinzler, B Vogelstein, and CA Iacobuzio-Donahue. Distant metastasis occurs late during the genetic evolution of pancreatic cancer. Nature, 467:1114–7, 2010.

[51] LR Yates and PJ Campbell. Evolution of the cancer genome. Nat Rev Genet, 13:795–806, 2012.

[52] J Zhang, J Fujimoto, J Zhang, DC Wedge, X Song, J Zhang, S Seth, CW Chow, Y Cao, C Gumbs, KA Gold, N Kalhor, L Little, H Mahadeshwar, C Moran, A Protopopov, H Sun, J Tang, X Wu, Y Ye, WN William, JJ Lee, JV Heymach, WK Hong, S Swisher, II Wistuba, and PA Futreal. Intratumor heterogeneity in localized lung adenocarcinomas delineated by multiregion sequencing. Science, 346:256–9, 2014.

[53] R Zhao and F Michor. Patterns of proliferative activity in the colonic crypt determine crypt stability and rates of somatic evolution. PLoS Comput Biol, 9:e1003082, 2013.

